# Nuclear localization of HD-Zip IV transcription factor GLABRA2 is driven by Importin α

**DOI:** 10.1101/2023.11.03.565550

**Authors:** Bilal Ahmad, Ruben Lerma-Reyes, Thiya Mukherjee, Hieu V. Nguyen, Audra L. Weber, Waltraud X. Schulze, Jeffrey R. Comer, Kathrin Schrick

## Abstract

GLABRA2 (GL2), a class IV homeodomain leucine-zipper (HD-Zip IV) transcription factor (TF) from *Arabidopsis*, is a developmental regulator of specialized cell types in the epidermis. GL2 contains a putative monopartite nuclear localization sequence (NLS) partially overlapping with its homeodomain (HD). We demonstrate that NLS deletion or alanine substitution of its basic residues (KRKRKK) affects nuclear localization and results in a loss-of-function phenotype. Fusion of the predicted NLS (GTNKRKRKKYHRH) to the fluorescent protein EYFP is sufficient for its nuclear localization in roots and trichomes. The functional NLS is evolutionarily conserved in a distinct subset of HD-Zip IV members including PROTODERMAL FACTOR2 (PDF2). Despite partial overlap of the NLS with the HD, genetic dissection of the NLS from PDF2 indicates that nuclear localization and DNA binding are separable functions. Affinity purification of GL2 from plant tissues followed by mass spectrometry-based proteomics identified Importin α (IMPα) isoforms as potential GL2 interactors. NLS structural prediction and molecular docking studies with IMPα-3 revealed major interacting residues. Split-ubiquitin cytosolic yeast two-hybrid assays suggest interaction between GL2 and four IMPα isoforms from *Arabidopsis.* Direct interactions were verified in vitro by co-immunoprecipitation with recombinant proteins. IMPα triple mutants (*impα- 1,2,3*) exhibit defects in EYFP:GL2 nuclear localization in trichomes but not in roots, consistent with tissue-specific and redundant functions of IMPα isoforms in *Arabidopsis*. Taken together, our findings provide mechanistic evidence for IMPα-dependent nuclear localization of GL2 and other HD-Zip IV TFs in plants.

**One sentence summary:** GLABRA2, a representative HD-Zip IV transcription factor from *Arabidopsis*, contains an evolutionarily conserved monopartite nuclear localization sequence that is recognized by Importin α for translocation to the nucleus, a process that is necessary for cell-type differentiation of the epidermis.

## INTRODUCTION

Class IV homeodomain leucine-zipper (HD-Zip IV) transcription factors (TFs) play a determinant role in regulation of plant epidermal and sub-epidermal differentiation (Nakamura et al., 2006; Chew et al., 2013; Schrick et al., 2023). HD-ZIP IV genes have been studied in various plant species, such as *Arabidopsis*, maize, rice, and cotton with the vast majority exhibiting epidermal-specific gene expression (Ingram et al., 2000; Nakamura et al., 2006; Depege-Fargeix et al., 2011). The founding member, *Arabidopsis* GLABRA2 (GL2), was first described for its role in trichome cell differentiation ∼30 years ago (Rerie et al., 1994). GL2 regulates various other developmental processes in the epidermis, including suppression of ectopic root hair formation (Di Cristina et al., 1996), seed coat mucilage biosynthesis (Western et al., 2000; Western et al., 2004), seed oil accumulation (Shen et al., 2006; Shi et al., 2012), and anthocyanin biosynthesis (Wang et al., 2015). Two other well-studied HD-Zip IV family members that are functionally redundant, ARABIDOPSIS THALIANA MERISTEM LAYER1 (ATML1) and PROTODERMAL FACTOR2 (PDF2), play a significant role in regulating embryonic and shoot epidermal development as well as in floral identity (Abe et al., 2003; Kamata et al., 2013; Ogawa et al., 2015; Iida and Takada, 2021).

It is well-established that TFs bind specific sites on the DNA to regulate gene expression after translocating to the nucleus. In the eukaryotes, proteins destined for subcellular localization to the nucleus are efficiently transported from their site of synthesis in the cytoplasm (Fu et al., 2018; Lu et al., 2021). Described as nucleocytoplasmic transport, this process enables the selective transport of proteins to mediate transcription, and other vital processes such as RNA processing, DNA replication and repair, and chromatin compaction (Liang and Clarke, 2001; Stewart, 2007; Marfori et al., 2011; Cautain et al., 2015; Kirby et al., 2015). This transport typically relies on the presence of a nuclear localization signal (NLS) in large proteins (>60 kDa), while smaller proteins (<40-60 kDa) may freely diffuse through the nuclear pore complex (NPC) (Cyert, 2001; Krebs et al., 2010).

The classical NLS, identified ∼40 years ago in SV40 T antigen and nucleoplasmin, is categorized as either a monopartite NLS, featuring a single stretch of positively charged amino acids (K-K/R-X-K/R, where X represents any residue), or a bipartite NLS, which consists of two interdependent stretches of basic residues separated by a variable spacer region ((K/R)(K/R)X10–12(K/R)3/5) (Dingwall et al., 1982; Kalderon et al., 1984; Wendler et al., 2004; Xu et al., 2010; Kirby et al., 2015; Soniat and Chook, 2015). One type of non-classical NLS known as the PY-NLS is characterized by central hydrophobic or basic motifs, followed by C-terminal R/K/H(X)2-5 PY motifs, and was structurally identified as an NLS in the human heterogeneous nuclear ribonucleoprotein A1 (hnRNP A1) (Lee et al., 2006).

Translocation through the NPC is mediated by karyopherins, commonly known as importins or exportins, depending on the transport direction (Meier and Brkljacic, 2009; Tamura and Hara-Nishimura, 2014). Importins are divided into distinct subfamilies representing two of the subunits of the import complex: Importin α (IMPα) and Importin β (IMPβ) (Tamura and Hara-Nishimura, 2014). Eukaryotic organisms contain multiple isoforms, and for example, the *Arabidopsis thaliana* genome encodes a total of 9 IMPα and 18 IMPβ subunits (Bhattacharjee et al., 2008; Liu et al., 2019). IMPα (Karyopherin α) was initially purified from erythrocytes (Adam and Geracet, 1991). Shortly thereafter, a fungal IMPα ortholog was cloned and characterized from yeast, based on its phenotype as a suppressor of temperature-sensitive RNA polymerase I mutations (SRP1) (Yano et al., 1992). A vertebrate ortholog was isolated from Xenopus eggs and referred to as Importin (Görlich et al., 1994). Thereafter, the first plant IMPα ortholog, an *Arabidopsis* protein, was reported and shown to recognize plant-derived NLS motifs and associate with the NPC (Hicks et al., 1996; Smith et al., 1997).

The steps of NLS-mediated translocation to the nucleus involve assembly of a cargo-carrier complex in the cytoplasm, translocation through the NPC, disassembly of the complex and release of the cargo in the nucleus, followed by recycling of the import machinery (Stewart, 2007). In one possible scenario, the IMPα subunit binds to a cargo protein via its NLS, forming a ternary complex with the IMPβ (Karyopherin β) subunit (Chook and Blobel, 2001; Liang and Clarke, 2001; Goldfarb et al., 2004; Lu et al., 2021). The IMPβ subunit subsequently engages in interactions with the NPC, and inside the nucleus it interacts with Ran-GTP followed by release of the cargo (Radu et al., 1995a; Radu et al., 1995b; Tamura and Hara-Nishimura, 2014). IMPβ may autonomously facilitate the nuclear translocation of cargo, acting as an intermediary, independently of IMPα (Chook and Süel, 2011). For instance, IMPβ directly associates with proteins containing the PY-NLS, subsequently transporting them into the nucleus, bypassing the requirement for IMPα (Gonzalez et al., 2021; Lu et al., 2021).

Structure-function studies of various *Arabidopsis* proteins provide clues on mechanisms underlying nuclear transport in plants. Sequence requirements for nuclear localization of cargos have been previously described by employing various heterologous transient expression systems. For example, an atypical NLS was identified in ASYMMETRIC LEAVES2-LIKE18/LATERAL ORGAN BOUNDARIES DOMAIN16 (ASL18/LBD16), a TF involved in lateral organ development, using a protoplast transfection assay to track the subcellular localization of GFP-tagged TFs (Kim and Kim, 2012). Mechanisms for nuclear import have been reported in a few cases that also relied on heterologous expression of cargo proteins. The nuclear import of Poly(ADP-Ribose) Polymerase 2 (PARP2) was investigated by expressing GFP-tagged cargos in *Nicotiana benthamiana*, and additional experiments suggested that *Arabidopsis* IMPα-2 mediates the nuclear translocation of PARP2 through a conserved monopartite NLS (Chen et al., 2018). In another study, the NLS motif requirements of the phosphoinositide kinase PHOSPHATIDYLINOSITOL 4-PHOSPHATE 5-KINASE 2 (PIP5K2) were deduced by expressing fluorescently-tagged proteins in onion epidermal cells, and protein-protein interaction assays suggested that IMPα isoforms bind to PIP5K2 to mediate nuclear import (Gerth et al., 2017).

In the present study, we sought to elucidate the nuclear localization mechanism of GL2 and other HD-Zip IV TF members from *Arabidopsis* within the native context of their expression in the plant. HD-Zip IV TFs are large ∼80 kDa proteins comprised of multiple domains including a Homeodomain (HD) DNA-binding domain, a Zipper-Loop- Zipper (ZLZ) dimerization domain, a Steroidogenic Acute Regulatory (StAR)-related lipid Transfer (START), and a START Adjacent Domain (STAD) (Schrick et al., 2023). The mechanism of nuclear localization of these key developmental regulators is not reported to date, although this information is critical for understanding how these TFs are controlled at the post-translational level. We investigated a predicted monopartite NLS, overlapping with the HD in GL2 and other family members from *Arabidopsis*.

Experiments with fluorescently-tagged GL2 expressed under its native promoter showed that the putative NLS is both necessary and sufficient for translocation to nuclei in root epidermal cells. Protein-protein interaction analysis using four independent approaches indicated that GL2 interacts with several IMPα isoforms, and additionally demonstrated that the NLS is required for this interaction. Consistent with the role of IMPα in the nuclear localization of GL2, *impα-1,2,3* triple mutants displayed trichome defects concomitant with nuclear import defects within trichome cells. We propose a model in which IMPα binds to the monopartite NLS in HD-Zip IV TFs to drive nuclear import, a mechanism that is critical for transcriptional regulation of epidermal development in plants.

## RESULTS

### Monopartite NLS is conserved in a subset of HD-Zip IV TFs in *Arabidopsis* and across the plants

Bioinformatic analysis of *Arabidopsis* GL2 revealed the presence of a 13-amino acid motif (96GTNKRKRKKYHRH108) that is predicted to function as an NLS (**Figure 1A**; **Supplemental Figure S1**; **Supplemental Table S1**). This NLS is categorized as a monopartite or “classical” NLS that contains a specific pattern of basic (K, R) residues overlapping with the N-terminal portion of the HD of GL2. Multiple sequence alignment and sequence analysis of all 16 HD-Zip IV TFs from *Arabidopsis* indicated that the predicted NLS is evolutionarily conserved in a subset comprising the majority (12 out of 16) family members including PDF2 and ATML1 (**Figure 1B and 1C**). In contrast, the other HD-Zip IV members (FWA, HDG8, HDG9, and HDG10) contain a putative bipartite NLS that varies in both sequence and position (**Supplemental Figure S1**). A phylogenetic analysis revealed that HDG8, HDG9, HDG10, and FWA form a distinct clade separate from other members of the family that contain a conserved monopartite NLS (**Figure 1B**).

**Figure 1.**
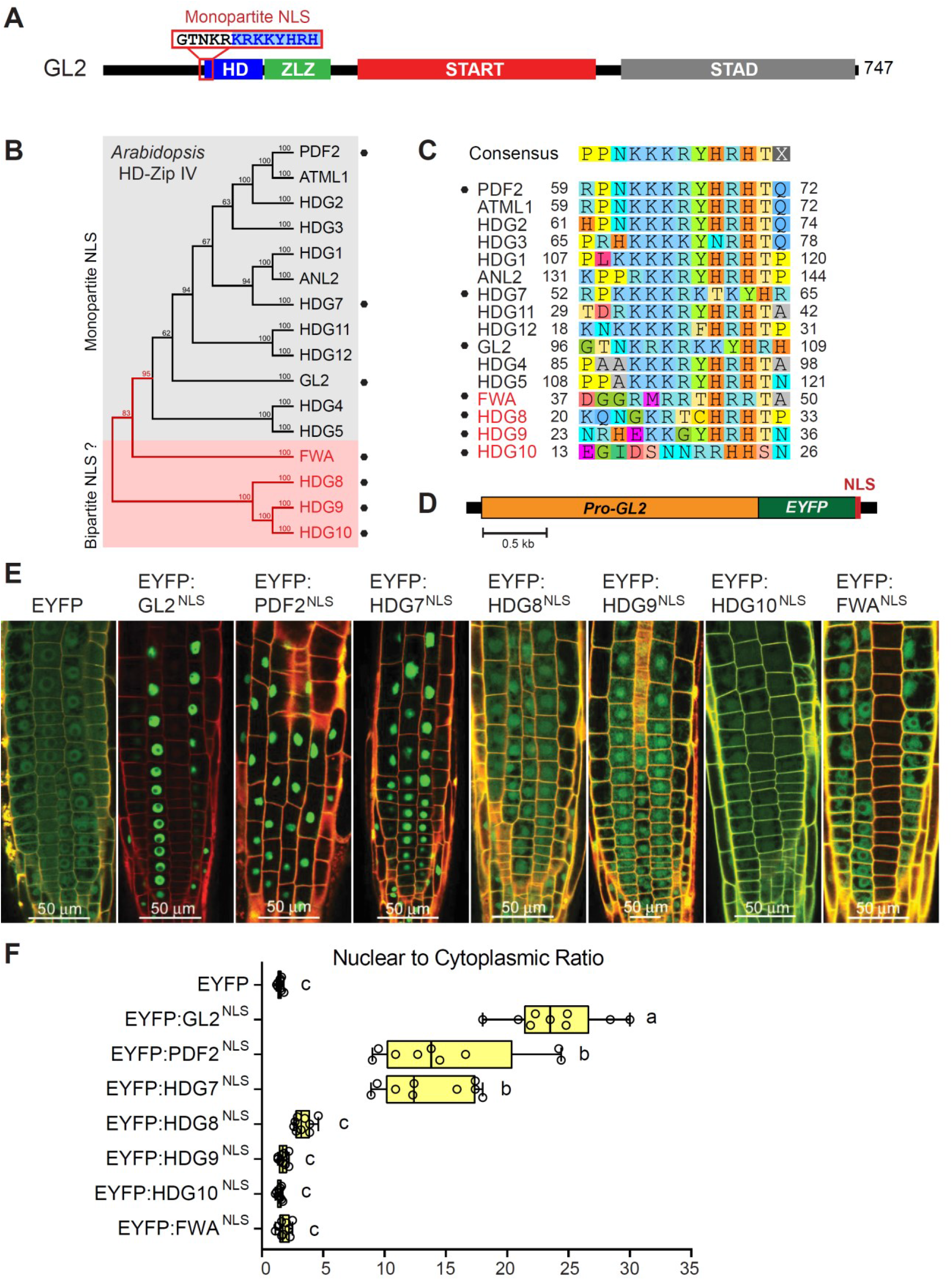
Predicted monopartite NLS from GL2, PDF2 and HDG7 is sufficient for nuclear localization of EYFP. (A) Domain configuration of GL2 indicating the position of the NLS, which partially overlaps with the HD. Homeodomain, HD; ZLZ, Zipper Loop Zipper; START, Steroidogenic Acute Regulatory (StAR) related lipid Transfer; STAD, START Adjacent Domain. (B) Phylogenetic tree illustrates sequence relationships among the 16 HD-Zip IV TFs from *Arabidopsis*. The tree was generated with full-length proteins using the unweighted pair group method with arithmetic mean (UPGMA) algorithm. Node numbers indicate consensus percentages with a bootstrap value of 1000. 12 out of 16 members, including GL2 and PDF2, contain a putative monopartite NLS, whereas the remaining members (FWA, HDG8, HDG9, and HDG10) contain a putative bipartite NLS. See also **Supplemental Figure S1** and **Supplemental Table S1**. (C) Predicted NLS motifs in the 16 HD-Zip IV TFs from *Arabidopsis*. Amino acid sequences and numbering are indicated. The alignment was determined by the 13 residue motif identified in GL2. (D) Schematic of the reporter construct in which the monopartite NLS from selected HD- Zip IV TFs was fused to the C-terminus of EYFP. The reporter gene was expressed under the *GL2* promoter (*GL2pro*) (E) Subcellular localization of EYFP alone in comparison to EYFP fusions to putative NLS sequences from the HD-Zip IV members given in (**B**). Roots from 3-4 day-old seedlings were imaged using confocal microscopy. EYFP fusions to predicted monopartite NLS of GL2, PDF2 or HDG7 result in nuclear localization of EYFP while aligned sequences from HDG8, HDG9, HG10, and FWA display both cytoplasmic and nuclear localization. (F) Quantification of nuclear to cytoplasmic ratio of EYFP pixels in three root epidermal cells from three seedlings of each line (n = 9). Representative images are shown in **(C)**. Box plots indicate minimum, maximum, 25^th^ and 75^th^ percentiles and median. Significant differences between genotypes were determined by one-way ANOVA, Tukey’s test, and are indicated by letters: p < 0.0001.

For the initial prediction of the NLS in GL2, we used cNLS Mapper, which applies an additivity-based motif scoring algorithm (Kosugi et al., 2009). NLStradamus, another NLS prediction tool having a hidden Markov model-based algorithm (Nguyen Ba et al., 2009), was utilized to search for NLS motifs in HD-Zip IV versus HD-Zip III family members from *Arabidopsis* (**Supplemental Table S1**). Both algorithms robustly predicted a monopartite NLS in the majority (12 out of 16) HD-Zip IV TF family members. In contrast, the five HD-Zip III TFs (AtHB8, CNA, PHB, PHV, REV) from *Arabidopsis* were predicted to contain a distinct NLS that partially overlaps with the C-terminus of their respective HD. Intriguingly, the NLS motifs predicted in HD-Zip IV versus HD-Zip III TFs both appear to be conserved in their respective *Spirogyra pratensis* orthologs from the charophycean green algae (**Supplemental Table S1**).

Multiple sequence alignments showed that the NLS from GL2 is highly conserved among GL2-like orthologs that are members of the dicots (**Supplemental Figure S2A**). In contrast, the specific NLS from PDF2 appears to be more deeply conserved across the land plants as well as in extant species from the charophycean green algae (**Supplemental Figure S2B**). For example, the HD-Zip IV ortholog from the freshwater green alga *Spirogyra pratensis*, predicted from sequencing data (Cooper and Delwiche, 2016) whose corresponding mRNA sequence we confirmed, contains a monopartite PDF2-like NLS (**Supplemental Table S1**).

### Conserved monopartite NLS from HD-Zip IV TFs is sufficient for nuclear localization of EYFP

To functionally test the predicted NLS from GL2 and other HD-Zip IV TFs, we systematically assessed the ability to import an enhanced yellow fluorescence protein (EYFP) reporter into the nuclei of *Arabidopsis* root epidermal cells. The 13-amino acid NLS or aligned amino acid stretch that overlaps with the HD, was taken from GL2, and other representative family members (PDF2, HDG7, HDG8, HDG9, HDG10, FWA) (**Figure 1C**). It was then fused to EYFP and expressed under the GL2 promoter (**Figure 1D**). The resulting *proGL2:EYFP:NLS* constructs were introduced into the *gl2* null mutant background to establish stable transgenic lines. A control construct lacking the NLS (*proGL2:EYFP*) was included. Examination of subcellular fluorescence distribution using confocal laser scanning microscopy demonstrated that the EYFP fused with the predicted NLS motif from GL2, PDF2, or HDG7 exhibited pronounced nuclear localization compared to the EYFP control. Conversely, the EYFP fusion with the analogous amino acid stretch from HDG8, HDG9, HDG10, and FWA failed to confer nuclear localization (**Figure 1E**). Quantitative analysis of fluorescence signals in the nucleus and cytoplasm revealed a higher nuclear-to-cytoplasmic ratio for EYFP fused with the NLS from GL2, PDF2, and HDG7, as compared to the EYFP control and EYFP fused to the corresponding 13-residue stretch from HDG8, HDG9, HDG10, and FWA (**Figure 1F**). These experiments provide evidence that the monopartite NLS conserved in GL2, PDF2, and HDG7, is sufficient nuclear localization of the EYFP reporter and that the sequence alone, rather than its position within the protein, is responsible for determining nuclear transport. As predicted from our bioinformatic analysis, our results indicate that the corresponding 13-residue stretches from HDG8, HDG9, HDG10, and FWA are not sufficient for nuclear import.

### Predicted NLS is necessary for nuclear import of GL2 and PDF2 in *Arabidopsis* roots

To investigate whether the NLS is required for nuclear transport of GL2, we initially tested two deletion mutants in comparison to wild type (**Figure 2A**). In the first mutant, the 61 residues encompassing the HD, including 9 out of 13 residues of the NLS, were deleted in frame. In the second mutant, only the 13-residue NLS motif of GL2 was deleted. Both mutant proteins lacking the HD or NLS (*gl2^ΔHD^* and *gl2^ΔNLS^*) and the wild-type *GL2* protein were expressed as EYFP-tagged proteins in the *gl2* null mutant background under the native GL2 promoter, as shown schematically in **Figure 2B**. The untransformed *gl2* null mutant lacked detectable EYFP fluorescence signal and served as a negative control. The wild-type transgenic line expressing *proGL2:EYFP:GL2* exhibited nuclear localization of EYFP:GL2 in distinct cell files of the root epidermis. In contrast, both the gl2^ΔHD^ and gl2^ΔNLS^ mutants exhibited indiscriminate subcellular localization throughout root epidermal cell files (**Figure 2C**). These findings provide evidence that the NLS is necessary for the nuclear localization of GL2.

**Figure 2.**
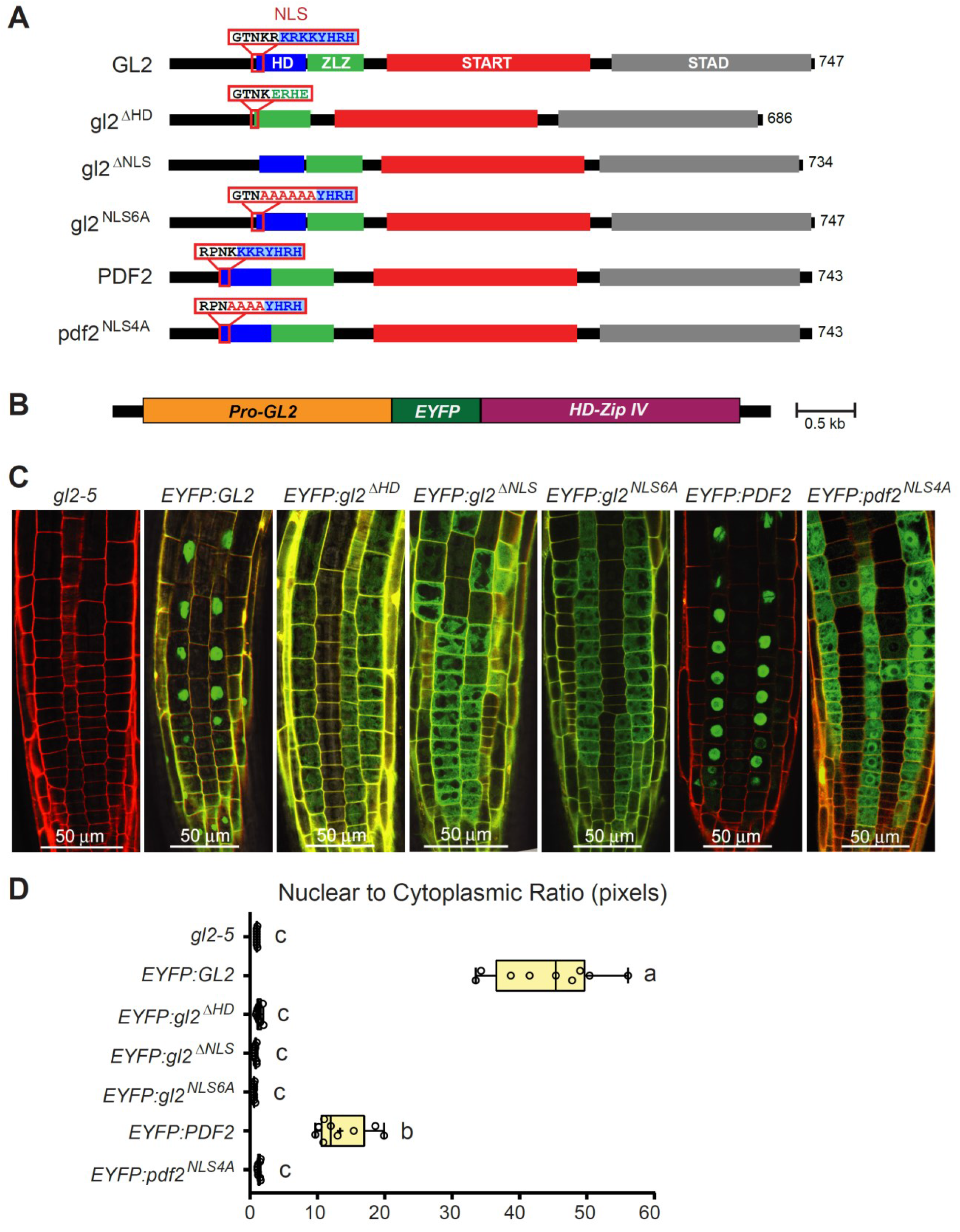
Predicted NLS is required for nuclear localization of EYFP-tagged GL2 and PDF2. **(A)** Domain configurations are illustrated in wild-type and mutant proteins. Position and sequence of the predicted NLS is indicated. HD, Homeodomain; ZLZ, Zipper Loop Zipper; START, Steroidogenic Acute Regulatory (StAR) related lipid Transfer; STAD, START Adjacent Domain. Amino acid lengths are shown on the right. **(B)** Schematic of the transgenic constructs for HD-Zip IV members GL2 and PDF2. The native *GL2* promoter (*proGL2*) drives the expression of EYFP-tagged TFs. **(C)** Subcellular localization of EYFP-tagged wild-type GL2 or PDF2 and corresponding NLS mutants. Roots from 3-4 day-old seedlings were imaged using confocal microscopy. The *gl2-5* background, in which the transgenes are expressed, lacks the EYFP signal. EYFP:GL2 and EYFP:PDF2 exhibit distinct nuclear localization compared to the mutant proteins lacking the putative NLS. **(D)** Quantification of nuclear to cytoplasmic ratio of EYFP pixels in three root epidermal cells from three seedlings of each line (n = 9). Representative images are shown in **(B)**. Box plots indicate minimum and maximum values, 25^th^ and 75^th^ percentiles and median. Significant differences between genotypes were determined by one-way ANOVA, Tukey’s test, and are indicated by letters: p < 0.0001.

We next constructed alanine substitution mutants to test whether the basic residues within the NLS motif are required for the nuclear localization of GL2 and another HD-Zip IV member, PDF2. The six basic residues (KRKRKK) from GL2 and the four basic residues (KKKR) from PDF2 were substituted with alanines, as shown in **Figure 2A**. The *proGL2:EYFP:gl2^NLS6A^*, *proGL2:EYFP:pdf2^NLS4A^*, and *proGL2:EYFP:PDF2* constructs were transformed into the *gl2* null mutant background to establish stable transgenic lines. Wild-type EYFP:GL2 and EYFP:PDF2 exhibited nuclear localization, whereas the corresponding EYFP:gl2^NLS6A^ and EYFP:pdf2^NLS4A^ mutants displayed indiscriminate subcellular localization (**Figure 2C**). Quantitative analysis of the data demonstrated a higher nuclear-to-cytoplasmic ratio of EYFP signal for each wild-type HD-Zip IV TF in comparison to the alanine substitution mutants (**Figure 2D**). These findings indicate that the basic residues within the NLS of GL2 and PDF2 are required for their nuclear import in cells of the root epidermis.

### GL2 NLS mutations display epidermal defects in leaves, roots and seeds, similar to those of the *gl2* null mutant

Our previous work demonstrated that deletion of the HD from GL2 results in a loss-of-function phenotype (Mukherjee et al., 2022). Here, we characterized the phenotypes of the transgenic lines expressing wild-type *GL2* in comparison to the NLS mutants, *gl2^ΔNLS^*, and *gl2^NLS6A^* in comparison to the *gl2^ΔHD^* mutant. While the *EYFP:GL2* expressing lines rescued the *gl2* null mutant and showed a trichome phenotype similar to that of wild type Columbia (Col), the NLS mutants resulted in glabrous leaves characteristic of the *gl2* null mutant control (**Figure 3A**). Quantification of the trichome phenotypes on first leaves of *gl2^ΔHD^*, *gl2^ΔNLS^*, and *gl2^NLS6A^* seedlings indicated branching defects that were indistinguishable from the *gl2* null mutant (**Figure 3B**). Similar to the subcellular localization patterns of the EYFP-tagged proteins in roots (**Figure 2**), we observed that only the wild-type EYFP:GL2 protein exhibited distinct nuclear localization in trichome cells (**Figure 3C**). Examination of primary roots from young seedlings and quantification of root hairs revealed that the NLS mutants display ectopic root hair formation that is indistinguishable from that observed for the *gl2* null mutant (**Figure 3D and 3E**). Similarly, we found that the NLS mutants exhibit *gl2* loss-of-function phenotypes in the production of seed coat mucilage (**Supplemental Figure S3**). Taken together, our mutant analysis demonstrates that the predicted monopartite NLS of GL2 is required for its proper function in trichomes, as well as in the root epidermis and seed coat cells.

**Figure 3.**
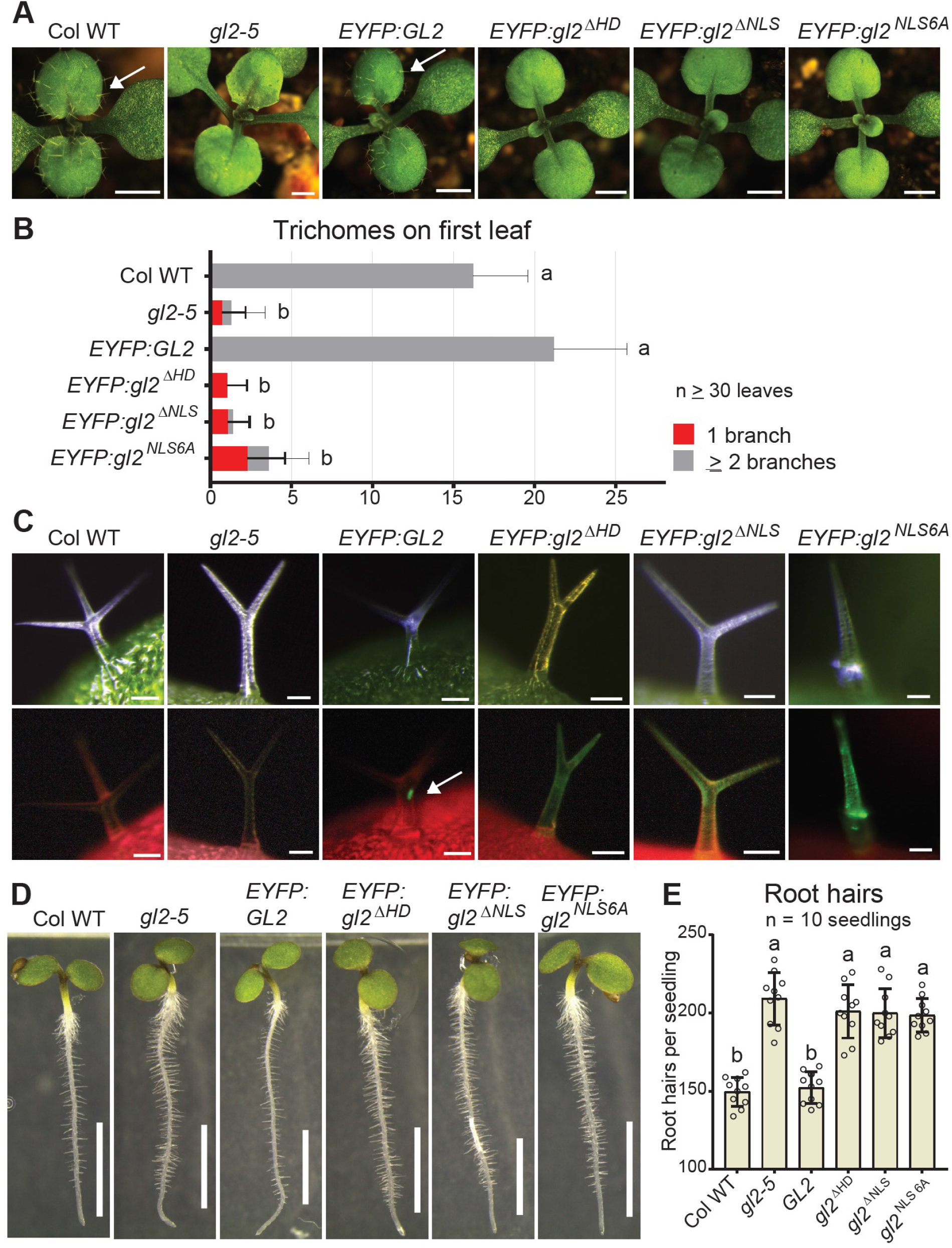
The *gl2* NLS mutants exhibit impaired nuclear localization along with defects in leaf trichomes and root epidermal cells. **(A)** Trichome phenotypes on the first leaves of *Arabidopsis* seedlings. Columbia wild type (Col WT) and *gl2-5* plants transgenic for *EYFP:GL2* exhibit trichomes on first leaves (arrows), while *gl2* NLS mutants (*gl2^ΔNLS^*, *gl2^NLS6A^*, *gl2^ΔHD^*) exhibit trichome defects similar to the *gl2-5* null mutant. Bar = 500 µm. **(B)** Quantification of trichomes and trichome branching patterns on first leaves. Representative images of seedlings are shown in **(A)**. Error bars indicate SD. Significant differences between genotypes were determined by one-way ANOVA, Tukey’s test, and are indicated by letters: p < 10^-5^. **(C)** Subcellular localization of EYFP-tagged wild-type GL2 and mutant gl2 proteins in trichomes. Imaging of leaf trichomes under white light (green, chlorophyll), with matching fluorescence images (red, chlorophyll). Nuclear localization is observed for EYFP:GL2 (arrow), but not for the EYFP-tagged mutants (*gl2^ΔHD^*, *gl2^ΔNLS^*, and *gl2^NLS6A^*). Col WT and *gl2-5* are negative controls for EYFP expression. Bar = 50 µm. **(D)** Root hair phenotypes of seedlings. Col WT and *EYFP:GL2* exhibit a normal root hair patterning, while *gl2* NLS mutants (*gl2^ΔNLS^*, *gl2^NLS6A^*, *gl2^ΔHD^*) display excessive root hair formation similar to the *gl2-5* null mutant. Bar = 1 mm. **(E)** Quantification of root hairs on primary roots of seedlings. Representative images of seedlings are shown in **(D)**. Error bars indicate SD. Significant differences between genotypes were determined by one-way ANOVA, Tukey’s test, and are indicated by letters: p < 0.0001.

### GL2 NLS deletion mutant is proficient in HD-Zip IV TF homodimerization

We considered the possibility that disruption of the NLS (or HD including part of the NLS) results in misfolded protein, and thus would not be able to be transported into the nucleus due to abnormal aggregation. Therefore, we investigated whether the gl2^ΔNLS^ mutant protein retains activity in other aspects of HD-Zip IV TF function. In a previous study, we established a homodimerization assay for GL2 using the yeast two-hybrid (Y2H) system, and we demonstrated that the gl2^ΔHD^ mutant protein is proficient in self-interaction (Mukherjee et al., 2022). The bait autoactivation assays revealed, that similar to wild-type GL2 and gl2^ΔHD^, the gl2^ΔNLS^ mutant protein does not display autoactivation and is therefore suitable for bait-prey interaction studies (**Supplemental Figure S4A**).

To assess the impact of NLS deletion on homodimerization, all possible bait and prey combinations of wild-type GL2 and mutant gl2^ΔNLS^ were evaluated. Like the wild-type GL2 and gl2^ΔHD^, gl2^ΔNLS^ exhibited self-interaction in homodimerization (mutant-mutant) and dimerization (wild type-mutant) assays (**Supplemental Figure S4B-C**). In our previous study we confirmed that the gl2^ΔHD^ mutant protein is properly expressed in yeast (Mukherjee et al., 2022). Here we used Western blotting to show that the gl2^ΔNLS^ mutant protein is expressed in yeast in a similar manner to the wild-type GL2 protein (**Supplemental Figure S4D**). The results indicate that the gl2^ΔNLS^ mutant protein is properly expressed and folding correctly, and furthermore that deletion of the NLS does not interfere with the homodimerization property of GL2.

### PDF2 nuclear localization and DNA binding are separable functions

We previously discovered that specific loss-of-function mutations in the HD affect DNA binding, despite nuclear localization of the corresponding mutant proteins when they are expressed as EYFP-tagged proteins (Mukherjee et al., 2022). A common feature of these specific HD mutants is that the predicted NLS is left intact, and therefore it is not surprising that they retain nuclear localization. Due to the striking amino acid overlap between the NLS and the HD, we were interested in investigating whether the disruption of the NLS has an impact on DNA binding. For these experiments, we focused on PDF2 as a representative HD-Zip IV TF, due to issues with GL2 in DNA-binding studies. To investigate whether the DNA binding and nuclear localization are separable functions, we utilized the pdf ^NLS4A^ mutant (described in **Figure 2**) alongside a stepwise series of additional alanine substitution mutants: pdf2^NLS1A^, pdf2^NLS2A^, and pdf2^NLS3A^ (**Figure 4A**). We constructed transgenic lines expressing the respective EYFP-tagged proteins to track nuclear localization in roots from young seedlings using confocal laser scanning microscopy (**Figure 4B**). Quantification of the nuclear to cytoplasmic ratios revealed that only the wild-type EYFP:PDF2 protein exhibited nuclear localization while the alanine substitution mutants pdf2^NLS4A^, and pdf2^NLS3A^ and pdf2^NLS2A^ showed indiscriminate subcellular localization (**Figure 4C**).

**Figure 4.**
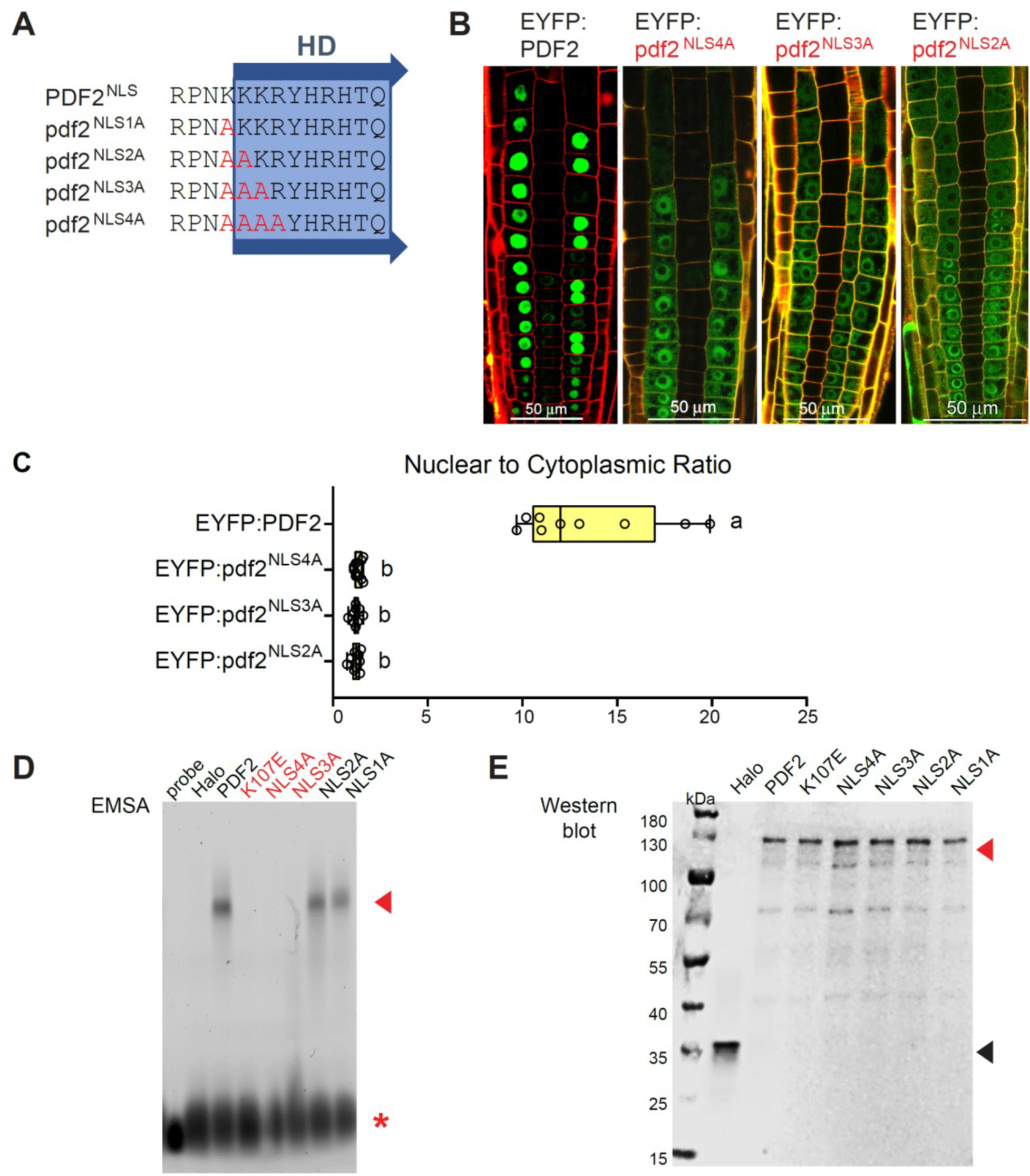
DNA binding and nuclear localization are separable functions. **(A)** Amino acid alignment showing the partial overlap between the putative monopartite NLS and the HD in PDF2. The wild-type sequence is shown in comparison to alanine substitution mutants that were experimentally tested for nuclear localization (**B**) and DNA binding (**C and D**). **(B)** Subcellular localization of EYFP-tagged wild-type GL2 or PDF2 and corresponding NLS mutants. Roots from 3-4 day-old seedlings were imaged using confocal microscopy. Wild-type EYFP:PDF2 exhibits nuclear localization in contrast to the alanine substitution mutants (pdf2^NLS4A^, pdf2^NLS3A^, pdf2^NLS2A^) which show both nuclear and cytoplasmic localization. **(C)** Quantification of nuclear to cytoplasmic ratio of EYFP pixels in three root epidermal cells from three seedlings of each line (n = 9). Representative images are shown in **(B)**. Box plots indicate minimum, maximum, 25^th^ and 75^th^ percentiles and median. Significant differences between genotypes were determined by one-way ANOVA, Tukey’s test, and are indicated by letters: p < 0.0001. **(D)** DNA binding of Halo-tagged wild type PDF2 and alanine substitution mutants to a fluorescein amidite (FAM)-labelled probe containing the L1 box in EMSA. Band shift (arrowhead) and free probe (asterisk) are indicated. Wild-type PDF2, pdf2^NLS2A^ and pdf2^NLS1A^ exhibit band shifts, while pdf2^NLS3A^ and pdf2^NLS4A^ are defective in DNA binding. An HD mutant (pdf2^K107E^) served as a negative control. The EMSA is representative of three replicate experiments. **(E)** Expression levels of input proteins used for the DNA binding experiments (**C**) are shown in Western blot with the Anti-HaloTag Ab. The Halo-tagged wild-type and mutant proteins (∼130 kDa) and Halo protein alone (∼33 kDa) are indicated (arrowheads). The Western blot is representative of three replicates.

A previously established electrophoretic mobility shifty assay (EMSA) with in vitro produced Halo-tagged proteins and a fluorescently-labeled probe (Mukherjee et al., 2022) was performed to test the DNA-binding activity of wild-type PDF2 and the NLS alanine substitution mutants. The HD loss-of-function mutant pdf2^K107E^, previously shown to be defective in DNA binding (Mukherjee et al., 2022), was included as a negative control. Our EMSA results revealed that the pdf2^NLS1A^ and pdf2^NLS2A^ mutants exhibit the expected band shift indicating their proficiency in DNA binding, similar to wild-type PDF2 (**Figure 4D**). In contrast, the pdf2^NLS3A^ and pdf2^NLS4A^ mutants, like pdf2^K107E^ and the vector control, failed to show a band shift, indicating defects in DNA binding (**Figure 4D**). Western blots showed comparable expression levels for the input proteins used in the EMSA, arguing against the possibility that the pdf2^NLS4A^ and pdf2^NLS3A^ mutants were poorly expressed (**Figure 4E**). Taken together, the experiments reveal that at least one NLS mutant, namely pdf2^NLS2A^, is defective in nuclear localization but retains DNA-binding ability, indicating that DNA binding and nuclear localization are distinct functions that can be separated.

### GL2 (NLS) interacts with IMPα in silico

Since classical monopartite NLS motifs are commonly found to be recognized by the IMPα subunit of the Importin complex, we considered five IMPα *Arabidopsis* isoforms (encoded by *IMPA-1, IMPA-2, IMPA-3, IMPA-4*, *IMPA-6*) as candidates for interaction with GL2 based on their tissue-specific expression patterns (Wirthmueller et al., 2013). Their corresponding amino acid sequences display high sequence similarity with rice IMPα-1 and mammalian and yeast IMPα orthologs (**Supplemental Figure S5A**). We next applied molecular docking, a computational approach for analyzing protein-peptide interactions (Wang et al., 2019), to investigate a possible interaction between the GL2- derived NLS and *Arabidopsis* IMPα-3, due to the availability of its crystal structure. The GL2 NLS (GTNKRKRKKYHRH) structure was predicted using PEP-FOLD (Lamiable et al., 2016), while the crystal structure of *Arabidopsis* IMPα-3 armadillo repeat domain (PDB ID: 4TNM) underwent refinement through homology modeling to address structural gaps (**Supplemental Figure S5B**). Ramachandran plot analysis of the IMPα- 3 refined model indicated that >97% of the residues reside in favored or fully allowed regions (**Supplemental Figure S6A**), and Qualitative Model ENergy Analysis (QMEAN) comparison to available structures in the Protein Data Bank (PDB) indicates a model of sharing geometrical features with known proteins (**Supplemental Figure S6B**).

Alignment of the IMPα-3 refined model with the template structure showed a root mean square deviation (RMSD) of 0.52, indicating that no significant structural changes occurred from resolving the gaps (**Supplemental Figure S5B**). The model is comprised of ten armadillo repeats, with the major NLS binding site at the N-terminus and the minor NLS binding site at the C-terminus (**Supplemental Figure S6C and S6D**).

Our protein-peptide docking simulations revealed that the GL2 NLS binds to the major binding site within the armadillo repeats of IMPα-3 (**Figure 5A**), consistent with previous findings that the classical monopartite NLS preferentially binds to the major site (Yang et al., 2010). Overall, the protein-peptide docking analysis revealed >30 specific interactions, encompassing both hydrogen bonding and electrostatic interactions, between GL2 NLS and IMPα-3, and highlighting the key residues involved in the binding interface (**Supplemental Table S2**). In particular, four charged residues of the GL2 NLS (K6,R7,K8,K9) play a central role in direct interactions with ten residues within the major binding site of IMPα-3 (**Figure 5A**). The IMPα-3 residues identified from computational docking with GL2 NLS are highly conserved in yeast, mouse, rice, and *Arabidopsis* (**Supplemental Figure S5A**), suggesting their functional importance. The presence of unique binding residues in our analysis, such as E106, could be attributed to several factors, such as variations in the lengths and surrounding sequences of basic residues, and/or structural differences between rice and *Arabidopsis* IMPα isoforms.

**Figure 5.**
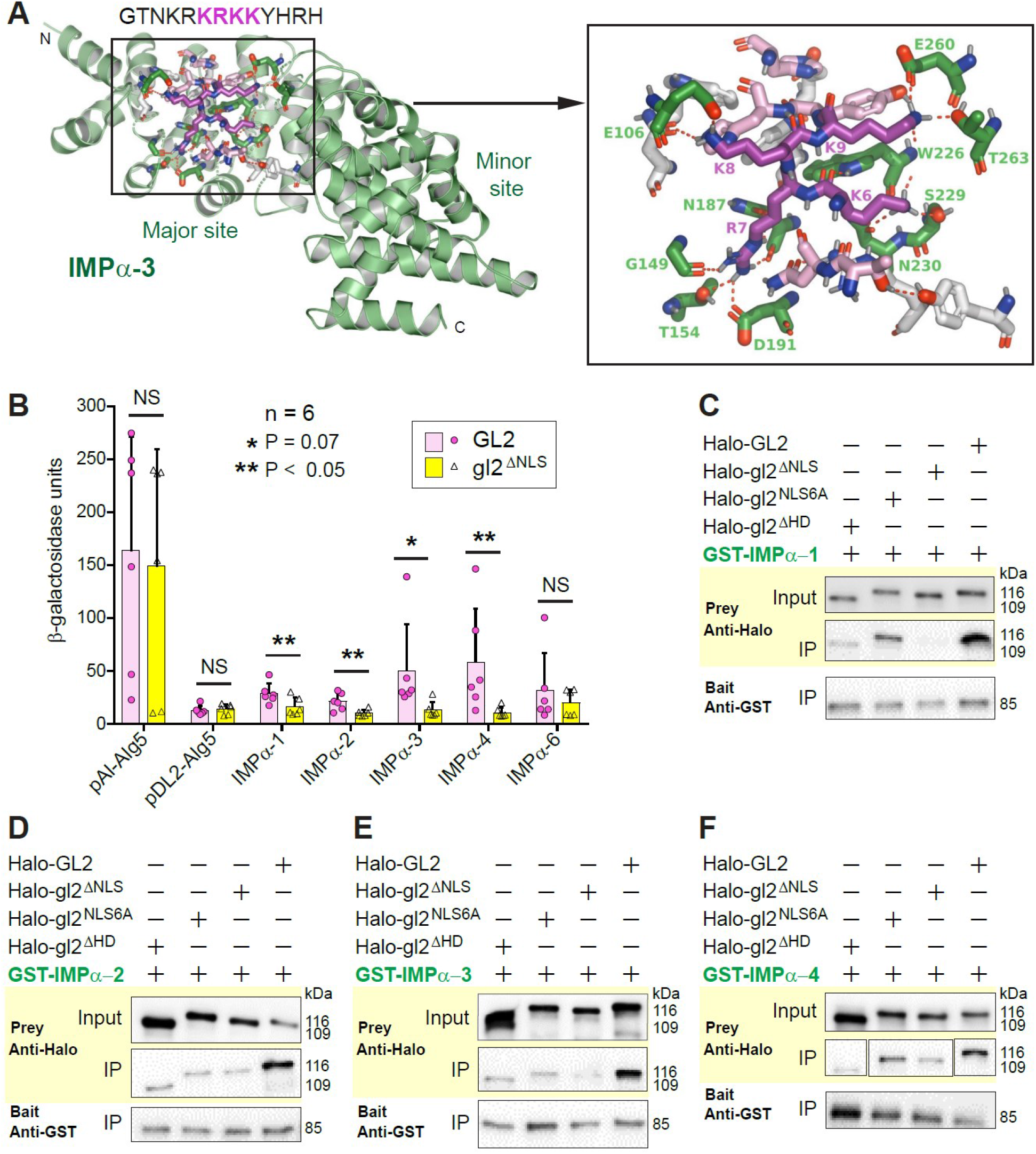
GL2 requires NLS to interact with IMPα. **(A)** Computational docking of the GL2 NLS (GTNKRKRKKYHRH) with *Arabidopsis* IMPα-3. Interacting residues of the NLS (magenta) and IMPα-3 (green) are indicated. Dotted lines depict direct molecular interactions between the NLS and IMPα-3. See **Supplemental Table S2**. **(B)** Split-ubiquitin-based cytosolic yeast two-hybrid (Y2H) quantitative β-galactosidase assay. The β-galactosidase units indicate *lacZ* reporter gene activities for the given bait (GL2 or gl2^ΔNLS^) and prey (IMPα) combinations. Interactions between wild-type GL2 (but not mutant gl2^ΔNLS^) and IMPα isoforms were detected for IMPα1, IMPα2, IMPα3 and IMPα4. *pAI-Alg5* and *pDL2-Alg5* served as positive and negative controls, respectively. Significant differences between GL2 and gl2^ΔNLS^ were determined by multiple *t*-tests: *p*- values are indicated. **(C-F)** Co-immunoprecipitation (Co-IP) experiments show that the GL2 NLS is important for direct physical interaction with IMPα. isoforms IMPα-1 **(C)**, IMPα-2 **(D)**, IMPα-3 **(E)**, and IMPα-4 **(F)**. GST-tagged IMPα isoforms served as baits while Halo-tagged wild-type GL2 or one of the NLS mutants (gl2^ΔNLS^, gl2^NLS6A^, gl2^ΔHD^) served as preys. Input indicates the initial protein amount used for the Co-IP. Input and immunoprecipitated (IP) proteins were detected in Western blots with Anti-Halo or Anti-GST Ab.

### NLS is required for in vivo interaction of GL2 with *Arabidopsis* IMPα isoforms in a yeast assay

To further investigate the physical interaction between GL2 and IMPα subunits, as well as the requirement of the NLS motif for this interaction, we employed a split-ubiquitin yeast two-hybrid assay (cytoY2H) (Möckli et al., 2007). In our experiments, wild-type GL2 and mutant gl2^ΔNLS^ were used as baits, while selected IMPα isoforms (IMPα-1, IMPα-2, IMPα-3, IMPα-4, and IMPα-6) served as preys. The interactions were quantified in terms of β-galactosidase units from activation of a *lacZ* reporter, and compared with positive and negative controls. From the cytoY2H assay results, IMPα-1, IMPα-2, IMPα- 3 and IMPα-4 show a significantly stronger interaction with wild-type GL2 as compared to with the NLS mutant gl2^ΔNLS^ (**Figure 5B**). These findings suggested that GL2 interacts physically with IMPα isoforms and that the NLS serves a crucial role in mediating this interaction.

### NLS is critical for interaction between GL2 and IMPα isoforms (IMPα-1, IMPα-2, IMPα-3, IMPα-4) in vitro

To further validate the results obtained from molecular docking and cytoY2H, we examined physical interactions between GL2 and IMPα isoforms using an in vitro co-immunoprecipitation (Co-IP) assay. We included several NLS mutants to serve as negative controls. In vitro transcription and translation of Halo-tagged wild-type GL2 and three NLS mutants (gl2^ΔHD^, gl2^ΔNLS^, gl2^NLS6A^) was employed for the production of the prey proteins. Recombinant expression in *E. coli* cells was used for the production of GST-tagged IMPα isoforms. The in vitro translated proteins were mixed with the bacterial cell extracts containing the respective IMPα isoforms, ensuring minimal or no perturbation to their native conformations. Subsequently, the protein mixtures were subjected to pull-down using anti-GST beads, followed by immunoblotting with the appropriate antibodies to detect the co-immunoprecipitated proteins. The results from the Co-IP assays revealed a strong binding between wild-type GL2 and each of the IMPα subunits that we tested (IMPα-1, IMPα-2, IMPα-3, and IMPα-4), while the gl2^ΔHD^, gl2^ΔNLS^, and gl2^NLS6A^ mutants exhibited significantly reduced binding (**Figure 5C-F**).

These findings provide further evidence, in addition to our in silico analysis and the cytoY2H experiments, underscoring the critical involvement of the NLS in mediating the physical interaction between GL2 and IMPα isoforms.

### GL2 pull-down uncovers IMPα-1, IMPα-2 and IMPα-6 as candidate interactors in ***Arabidopsis* plants**

The interaction of GL2 with members of the IMPα family was additionally confirmed by pull-downs from hydroponically grown *Arabidopsis* plants using EYFP:GL2 as a bait (**Supplemental Figure S7; Supplemental Data Set 1**). Interacting proteins were identified by mass spectrometry, and the intensity based absolute quantification (iBAQ) algorithm was used to determine the abundance of identified proteins within the GL2 interactome. A total of 2484 proteins were identified, including the bait protein GL2.

Details about the GL2 interactome will be provided in a separate publication. Relevant to the present work, the IMPα-1, IMPα-2 and IMPα-6 isoforms from *Arabidopsis* were found among 12 proteins with diverse functions in nuclear targeting (**Supplemental Figure S7**), consistent with our other protein-protein interaction studies.

### The *impα-1,2,3* mutant exhibits impaired GL2 nuclear localization in trichomes

Next, we took a genetic approach to assess the impact of IMPα function on GL2 nuclear localization. A previously characterized *impα-1,2,3* triple mutant (Chen et al., 2020) was crossed with the *EYFP:GL2* transgenic line to study the subcellular localization of GL2 in wild-type versus mutant backgrounds. Strikingly, fluorescence microscopy of trichome cells revealed that *impα-1,2,3* mutants exhibit aberrant nuclear localization of EYFP:GL2 (**Figure 6A**). Quantitative analysis indicated that approximately 16% of trichomes displayed a non-nuclear localization of EYFP:GL2, while 18% exhibited both nuclear and cytoplasmic localization (**Figure 6B**). In contrast, no nuclear localization defects were seen in control EYFP:GL2 lines that were wild type for the IMPα genes.

**Figure 6.**
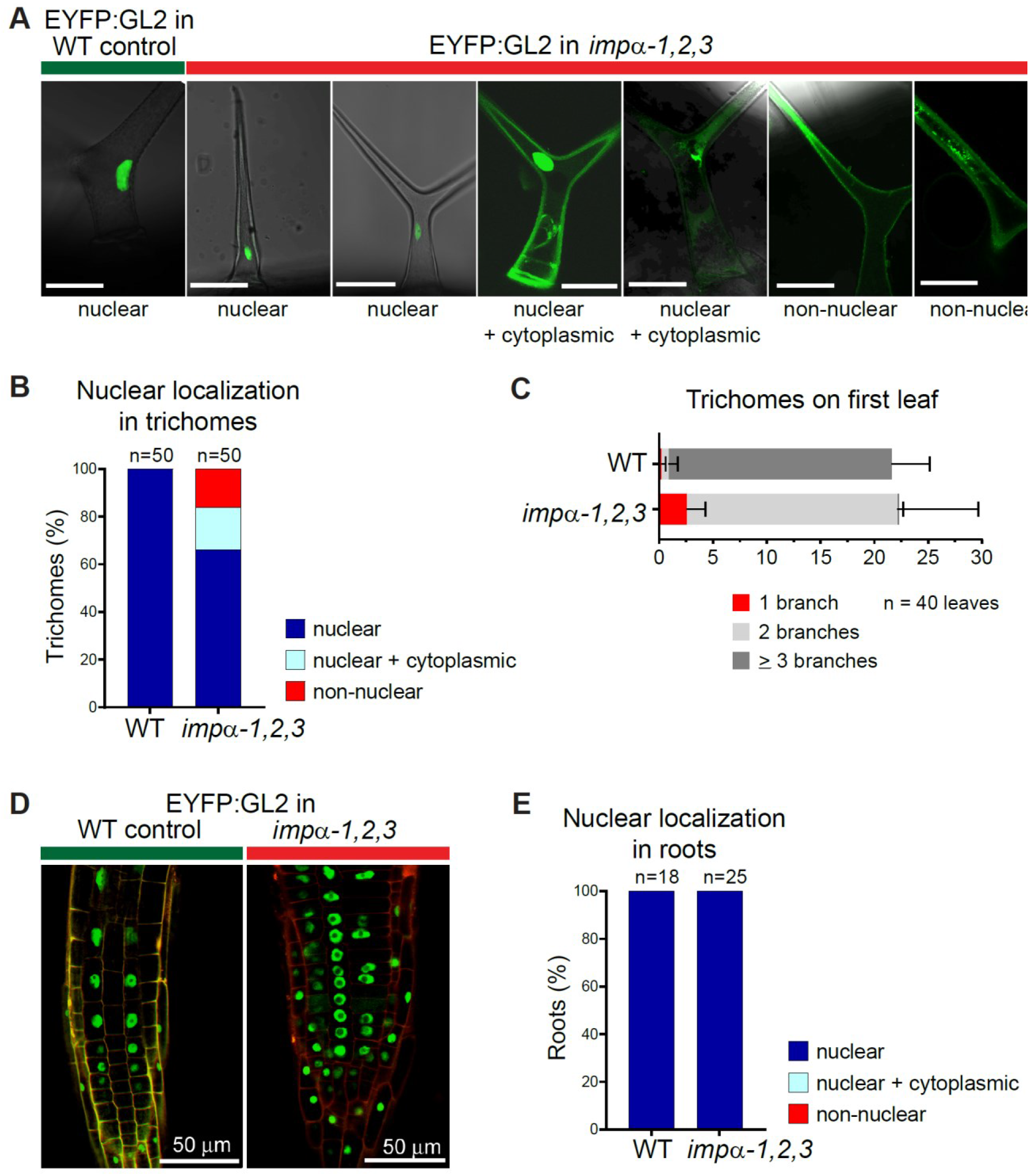
IMPα triple mutant (*impα-1,2,3*) is defective in nuclear localization of EYFP:GL2 in trichomes but not in roots. **(A)** Wild-type *IMPα* seedlings display nuclear localization of EYFP:GL2 in trichomes on first leaves. In contrast, *impα-1,2,3* seedlings expressing EYFP:GL2 are characterized by a heterogeneous population of trichome cells exhibiting nuclear, nuclear and cytoplasmic, and non-nuclear localization of EYFP:GL2. Bar = 50 µm. **(B)** Quantification of subcellular distribution of EYFP:GL2 in trichomes from wild-type *IMPα* and mutant *impα-1,2,3* seedlings. Representative images of nuclear, nuclear and cytoplasmic, and non-nuclear expression patterns are shown in **(A)**. A total of n=50 trichomes were assessed for each genotype. **(C)** Quantification of trichomes numbers and branches on first leaves from wild-type *IMPα* and mutant *impα-1,2,3* seedlings expressing EYFP:GL2. Trichomes from n=40 leaves were quantified for each genotype. **(D)** Confocal microscopy of primary roots from wild-type *IMPα* and mutant *impα-1,2,3* seedlings. Both genotypes display nuclear localization of EYFP:GL2 in roots. Bar = 50 µm. **(E)** Quantification of EYFP:GL2 subcellular localization in roots from wild-type *IMPα* and mutant *impα-1,2,3* seedlings. At least n=18 roots were assessed for each genotype.

We additionally detected trichome differentiation defects in the *impα-1,2,3* mutant seedlings (**Figure 6C**), as expected from the observed GL2 nuclear localization defects.

In contrast, in primary root tissues, no defects in nuclear localization of EYFP:GL2 were observed in *impα-1,2,3* mutant and control lines (**Figure 6D and 6E**). This suggests a functional redundancy of IMPα isoforms, which may compensate for nuclear localization in the absence of *IMPα-1, IMPα-2,* and *IMPα-3*. Our findings are consistent with previously reported tissue and cell-type specific gene expression profiles of the nine *Arabidopsis* IMPα isoforms (Wirthmueller et al., 2013), in particular IMPα-4 and IMPα-6 (**Supplemental Figure S8**). Since GL2 interacts with IMPα-1, 2, 3, and 4 in Co-IP (**Figure 5C-F**), it is possible that IMPα-4, which was shown to be expressed in the root, might contribute to GL2 nuclear localization in roots.

## DISCUSSION

### Evolutionary divergence of the NLS among HD-Zip IV TF family members

The overall goal of the present study was to decipher the nuclear localization mechanism of the HD-Zip IV TFs, a family of transcriptional regulators that control gene expression networks underlying epidermal development in plants (Nakamura et al., 2006; Schrick et al., 2023). Based on the data presented herein, we propose a model for an IMPα-dependent nuclear import mechanism governing the subcellular localization of HD-Zip IV TFs, taking GL2 as an example (**Figure 7**). Our initial bioinformatic analysis identified a predicted classical monopartite NLS in a subset of 12 out of 16 HD-Zip IV family members in *Arabidopsis* (**Supplemental Figure S1**). This finding is surprising considering the high degree of sequence conservation of the HD DNA- binding domain within these proteins (Mukherjee et al., 2022). Nonetheless, the GL2- like NLS is conserved across GL2 orthologs across the dicots, and the PDF-like NLS occurs in HD-Zip IV TFs across the plant kingdom including mosses and charophycean green algae, implying a striking evolutionary conservation of this NLS motif, albeit in a subset of family members (**Supplemental Figure S2**).

**Figure 7.**
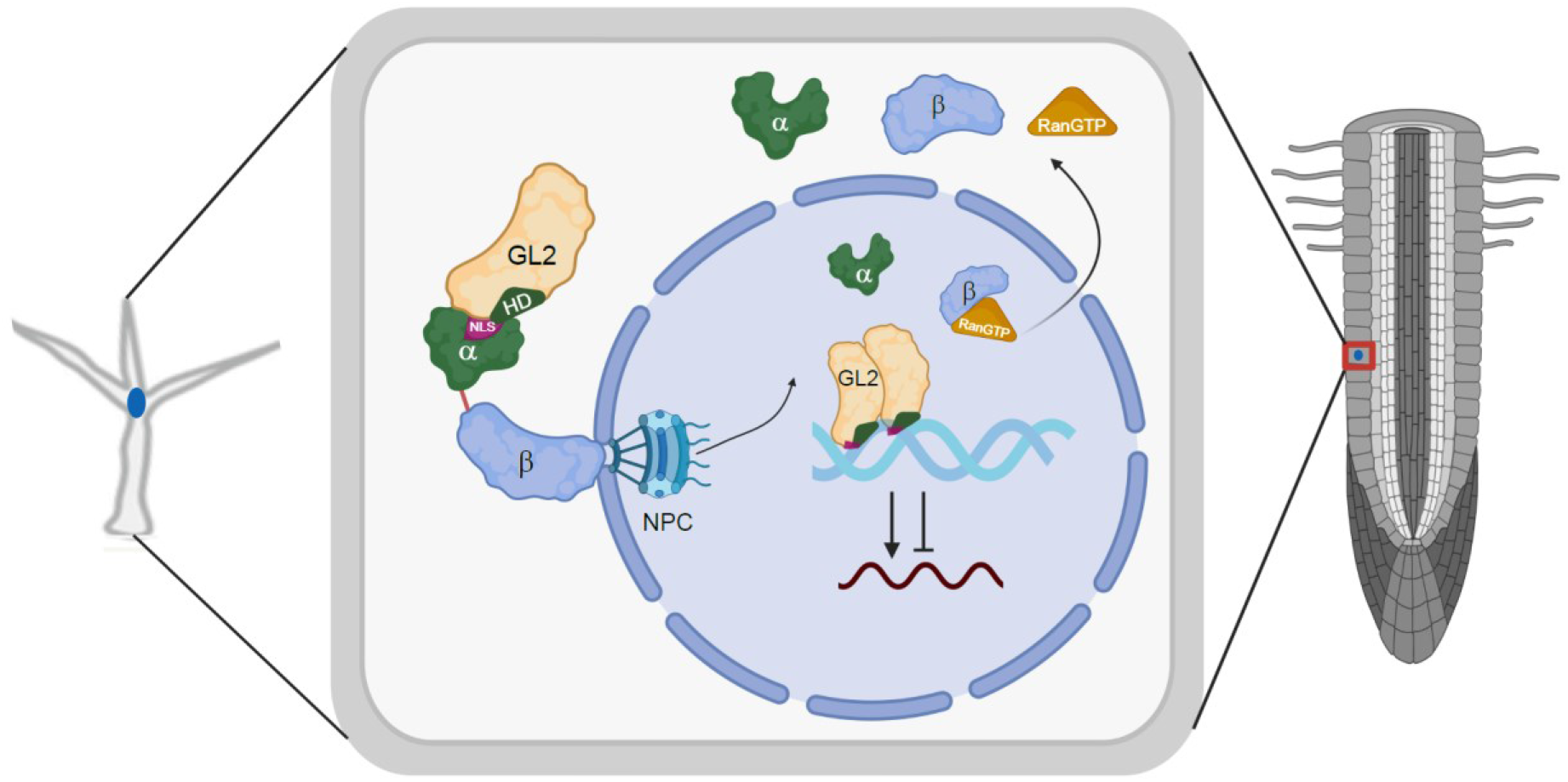
Model for nuclear import mechanism of GL2. IMPα isoforms facilitate GL2 nuclear localization in trichomes (left) and in the root epidermis (right). The monopartite NLS (Magenta), which partially overlaps with the HD (dark green), binds to IMPα (green), which in turn binds to IMPβ (blue). The GL2- IMPα/β complex interacts with nucleoporins of the nuclear pore complex (NPC) to facilitate nuclear transport of the trimer. In the nucleus, Ran GTP (orange) binds to IMPβ, resulting in the dissociation of the complex and GL2 release. Subsequently, GL2 forms a homodimer and binds to the DNA to positively or negatively regulate gene expression of its transcriptional targets.

The presence of a PDF2-like NLS in the extant freshwater green algae *Spirogyra pratensis* suggests that the four divergent HD-Zip IV family members (HDG8, HDG9, HDG10, FWA) that lack a functional NLS, at the same position overlapping with the HD, have lost this motif during the course of evolution. Consistent with our findings, a GFP fusion to the N-terminal part of FWA, comprised of the putative NLS and the HD DNA binding domain, displayed both nuclear and cytoplasmic localization in the female gametophyte and developing endosperm (Kinoshita et al., 2004), similar to what we observed for the EYFP:FWA^NLS^ reporter in young roots (**Figure 1E-F**). Intriguingly, the full-length FWA protein tagged with GFP similarly exhibits both nuclear and cytoplasmic localization (Kinoshita et al., 2004), but since the data were not quantified we cannot be sure about the level of indiscriminate subcellular localization. Our quantitative data indicate that the non-functional NLS of FWA exhibits indiscriminate subcellular localization, with a significantly lower nuclear-to-cytoplasmic ratio compared to the functional monopartite NLS of GL2, PDF2, and HDG7 (**Figure 1F**). It is noteworthy that we expressed EYFP:FWA^NLS^ under the GL2 promoter instead of the FWA promoter. In its native context, FWA may interact with IMPα isoforms (IMPα-5, 7, 8, or 9) that are not highly expressed in root epidermal cells (Wirthmueller et al., 2013). Moreover, in our study we used a short 13-residue stretch instead of the full-length protein. Perhaps other sequences within the full-length FWA protein mediate interactions with importins. Bioinformatic analysis based on a hidden Markov model (Nguyen Ba et al., 2009) predicts two NLS motifs in FWA: one at the N-terminus (which failed to behave as an NLS in our study) and the other at the C-terminus (which was not tested here) (**Supplemental Table S1**). An additivity-based motif scoring algorithm (Kosugi et al., 2009) predicts that FWA contains a bipartite NLS within the START domain (**Supplemental Figure S1**). It is possible that FWA, HDG8, HDG9 and HDG10 have evolved a novel mechanism for their nuclear localization, or alternatively, they possess critical functions outside the nucleus.

### Monopartite NLS of GL2 and PDF2 is critical for nuclear localization but not for homodimerization or DNA binding

By definition, a functional NLS must be both necessary and sufficient for the nuclear localization of a cargo protein (Lange et al., 2007; Lu et al., 2021). Our findings fulfill both requirements in the case of the monopartite NLS motifs found in GL2 and PDF2. We showed that fusing the corresponding 13-residue sequences from GL2 and PDF2 to an EYFP reporter results in the nuclear localization of the fluorescent protein EYFP (**Figure 1**). Deletion or substitution of basic residues in GL2 (KRKRKK) and PDF2 (KKKR) results in the loss of nuclear localization, further highlighting the essential role of the NLS in directing these HD-Zip IV TFs to the nucleus, and the GL2 NLS is required for its function in epidermal development (**Figures 2 and 3; Supplemental Figure S3**).

Homodimer formation by GL2 is critical for its activity, and it was previously shown that mutations in the START domain of GL2 impair the ability to form a homodimer although these mutations do not affect nuclear localization (Schrick et al., 2014; Mukherjee et al., 2022). In the present study, we utilized Y2H assays to demonstrate that the dimerization property of GL2 remains unaffected even in the absence of the NLS, such as in *gl2^ΔNLS^* mutants (**Supplemental Figure S4**).

Since the NLS of GL2 and PDF2 overlaps with the HD, we addressed the relationship between nuclear localization and DNA-binding, which is mediated by the HD. Our previous work indicated that a missense mutation in the HD (*pdf2^K107E^*) disrupts DNA binding, but it does interfere with nuclear localization of the mutant protein (Mukherjee et al., 2022). The analogous mutation in GL2 (*gl2^K146E^*) along with deletions affecting the HD but not the NLS, also result in nonfunctional protein that is nonetheless nuclear localized (Mukherjee et al., 2022). In the present study our EMSA experiments indicated that the alanine substitution mutants *pdf2^NLS2A^* and *pdf2^NLS1A^* retain the ability to bind DNA, despite the *pdf2^NLS2A^* mutant being defective in nuclear localization (**Figure 4**). Altogether, our data indicate that DNA binding and nuclear localization are two distinct functions, and that NLS mutants undergo proper protein folding but are specifically defective in nuclear localization.

### The NLS is required for GL2 recognition and binding to IMPα isoforms

In our cytoY2H experiments, the interaction between GL2 and *Arabidopsis* IMPα isoforms appeared weak, albeit significant differences were observed for GL2 versus g2^ΔNLS^ (**Figure 5B**), indicating the critical importance of the NLS. Previous structural studies confirmed that plant-derived NLS motifs can bind to yeast and mammalian IMPα orthologs (Chang et al., 2012). Thus, the weak interaction that we observed in the yeast system may be attributed to competition from yeast NLS-containing proteins and/or displacement of GL2 by yeast IMPα.

The Co-IP results corroborated the requirement of the GL2 NLS for the interaction with *Arabidopsis* IMPα isoforms (**Figure 5C-F**). Weak binding observed between NLS mutants and some of the IMPα isoforms can be attributed to the presence of other basic residues that can trigger the interaction during Co-IP. Considering our in planta data, these interactions alone are inadequate for the nuclear localization of the mutant proteins (**Figure 2**). The cytoY2H, Co-IP, and pull-down experiments collectively indicate that GL2 can interact with IMPα isoforms. However, we do not know whether GL2 prefers a specific IMPα isoform when multiple isoforms are present. Preferences could be influenced by various factors, including the expression levels of IMPα isoforms, their availability due to tissue-specific expression, post-translational regulation, and/or competition among cargo proteins (Wirthmueller et al., 2015). Our data are consistent with previous observations that IMPα family members exhibit preferences for specific NLS cargo, although some functional redundancy exists (Köhler et al., 1999; Rebane et al., 2004; Fagerlund et al., 2005; Kelley et al., 2010; Chen et al., 2018; Lüdke et al., 2021).

### Regulation of HD-Zip IV TF nuclear transport

While our study showcases the fundamental mechanism governing the nuclear translocation of HD-Zip IV TFs (**Figure 7**), it is intriguing to consider additional factors that might influence subcellular localization. The process of nuclear localization is highly dynamic and may be subject to spatiotemporal control (Saez et al., 2011; Yumerefendi et al., 2015). For example, nuclear localization of Phytochrome B is light-dependent (Chen et al., 2005). Post-translational modifications, such as phosphorylation and acetylation are additionally known to play roles in the regulation of nuclear localization (Poon and Jans, 2005; Yumerefendi et al., 2015; Su et al., 2018). During Influenza A virus infection, phosphorylation and dephosphorylation of key residues on the viral nucleoprotein impact binding affinity with IMPα isoforms, thereby controlling its entry into nuclei (Li et al., 2015; Zheng et al., 2015). Intra- or intermolecular interactions leading to conformational changes may additionally serve as switches in controlling nuclear localization (Poon and Jans, 2005; Bai et al., 2017; Liu et al., 2018). Another consideration is that the regulation of nuclear import of specific cargos may be influenced by post-translational modifications on IMPα itself. Palmitoylation of Xenopus IMPα directs it to the plasma membrane (PM), modulating its availability to NLS-containing cargos (Brownlee and Heald, 2019). In another example, the binding of IMPα to heparin within heparan sulfate proteoglycans targets it to the PM (Ma et al., 2021), making it unable to act in nuclear import.

Intriguingly, the HD-Zip IV TFs GL2 and ATML1 are both reported to exhibit differential localization between the nucleus and cytoplasm according to developmental stage (Szymanski et al., 1998; Iida et al., 2019). Immunocytochemical studies with anti- GL2 antibodies indicated that in subepidermal cells of young leaves, GL2 is found throughout cytoplasm as well as in nuclei (Szymanski et al., 1998). However, in trichomes at later stages, GL2 exhibits strong nuclear localization. More recent studies with GFP-tagged proteins in *Arabidopsis* embryos indicate that ATML1 is predominantly nuclear localized until the eight-cell stage, but at the 32-cell stage, it exhibits both nuclear and cytoplasmic localization (Iida et al., 2019). These observations suggest dynamic subcellular distributions of HD-Zip IV TFs, implying complex regulatory mechanisms controlling nuclear localization. Future studies addressing novel post-translational mechanisms underlying the functions of HD-Zip IV TFs and their interactions with IMPα isoforms will help in our understanding of the complexities governing their fascinating activities and movements within cells.

## MATERIALS AND METHODS

### Plant material, growth conditions and *Arabidopsis* transformation

*Arabidopsis* plants were of the Columbia (Col) ecotype. The *gl2-5* null mutant was described previously (Khosla et al., 2014). The *impα-1,2,3* triple mutant (Salk_001092; Salk_099707; Salk_025919) was provided by Jae Bok Heo and is described in (Chen et al., 2020). The *impα-1,2,3* mutant was crossed to the previously described *proGL2:EYFP:GL2* transgenic line (Khosla et al., 2014) and F2 progeny were genotyped for presence of the corresponding *impα* alleles using primers listed in **Supplemental Table S3**. Plants were stratified at 4°C for 4-5 days and grown on soil containing Metro-Mix 380, vermiculite, and perlite (Hummert International, Topeka, KS) in a 4:2:1 ratio at 23°C under continuous light. *Arabidopsis gl2-5* plants were transformed with *Agrobacterium tumefaciens* GV3101 (pMP90) using floral dip (Clough and Bent, 1998). T1 transformants were selected on 20 μg/ml Hygromycin B, followed by transfer of at least 20 seedlings per construct to soil. T1 plants were screened for EYFP expression, and a T2 segregation ratio of 3:1 Hygromycin B resistance. For each construct, 2-4 independent transformants were initially characterized, and at least one representative homozygous T3 line was genotyped by PCR and verified by sequencing.

### Binary Vector and plasmid construction

The EYFP reporter constructs for nuclear localization were created by bridging the linearized *proGL2:EYFP* SR54 binary vector lacking the GL2 cDNA with ssDNA oligos using the NEBuilder HiFi DNA Assembly Kit (New England Biolabs) and oligonucleotides given in **Supplemental Table S3**. The SR54 binary vectors carrying *proGL2:EYFP:GL2* and *proGL2:EYFP:gl2^ΔHD^* were described previously (Schrick et al., 2014; Mukherjee et al., 2022). The *proGL2:EYFP:gl2^ΔNLS^* construct was created using the Q5 Site-Directed Mutagenesis Kit (New England Biolabs), taking wild-type GL2 cDNA in pENTR/D-TOPO as a template, along with the primers listed in **Supplemental Table S3**. The SR54 binary vector was restriction digested with *Sal*I and *Kpn*I. Q5 High Fidelity DNA Polymerase (New England Biolabs) was used to amplify the mutant versions from their respective pENTR/D-TOPO vectors, using primers listed in **Supplemental Table S3**. NEBuilder HiFi DNA Assembly was employed to assemble the cDNA sequences into the linearized SR54 binary vector. To generate alanine substitutions within the KRKRKK NLS motif, GL2 was amplified in two fragments (294 bp and 1932 bp) using the primers (eYFP-GL2_F and gl2_NLS_KRK99AAA-R; and gl2_NLS_KRK99AAA_F and p1300-GL2-R) with the relevant mutations inserted in their 5’ ends. NEBuilder Hifi DNA Assembly was conducted with *Sal*I and *Kpn*I linearized SR54 vector. This method was similarly used to construct *proGL2:EYFP:pdf2^NLS4A^* with the EYFP-PDF2_F, pdf2_NLS_KRK62AAA-R and pdf2_NLS_KRK62AAA-R/p1300- PDF2-R primers. The *pdf2^NLS3A^*, *pdf2^NLS2A^* and *pdf2^NLS1A^* binary constructs were also assembled with the SR54 binary vector using the NEBuilder Hifi DNA Assembly Kit.

Plasmids for the cytoY2H studies were constructed as follows: The GL2 cDNA was cloned into the pBT3-C-OST4 plasmid (Gerth et al., 2017) using restriction digestion of the vector with *Sma*I followed by NEBuilder Hifi DNA Assembly using primers listed in **Supplemental Table S3**. pRR3-N Prey plasmids containing cDNAs for the *Arabidopsis* isoforms IMPα-1, IMPα-2, IMPα-3, IMPα-4, and IMPα-6 were a gift from Mareike Heilmann. The plasmids containing N-terminally Halo-tagged *GL2*, *gl2^ΔNLS13^*, *gl2^NLS6A^*, *pdf2^NLS4A^, pdf2^NLS3A^, pdf2^NLS2A^,* and *pdf2^NLS1A^* were constructed using Q5 Site-Directed Mutagenesis (New England Biolabs) to mutagenize *GL2* and *PDF2* in their pENTR/D-TOPO vectors with oligonucleotide primers given in **Supplemental Table S3**.

Thereafter, the wild-type and mutant versions were inserted in the pIX-HALO destination vector using the Gateway LR Clonase II Enzyme Mix (Invitrogen). Positive clones were screened by colony PCR and confirmed by Sanger sequencing. To construct the GST:IMPα fusions for recombinant expression in *E. coli*, IMPα isoforms were cloned in pGEX-6P-1 using the NEBuilder Hifi DNA Assembly Kit. The pGEX-6P-1 vector was digested with *BamH*I and *Sal*I, and the IMPα cDNA sequences were amplified from the corresponding pPR3-N plasmids using Q5 High Fidelity DNA Polymerase and primers listed in **Supplemental Table S3**.

### Phylogenetic analysis

Multiple sequence alignment of the full-length proteins corresponding to the 16 HD-Zip IV family members from *Arabidopsis* was performed using MUSCLE (Edgar, 2004) in MEGA11 (Tamura et al., 2021) with default settings. The evolutionary distances were computed using the Poisson correction. Ambiguous positions in each sequence pair were removed using the pairwise deletion option. Subsequently, a phylogenetic tree was constructed using the Unweighted Pair Group Method with Arithmetic Mean (UPGMA) method with 1000 bootstrap replicates.

### Bioinformatic prediction of NLS motifs

To initially predict the NLS in GL2 and other HD-Zip IV TF members, we used an web-based computer program (cNLS Mapper, http://nls-mapper.iab.keio.ac.jp/), which calculates IMPα/β-pathway NLS probability by applying an additivity-based motif scoring algorithm (Kosugi et al., 2009). At the time of submission, cNLS Mapper was no longer publicly available. Therefore we utilized another tool to validate the classical NLS predictions, NLStradamus (NovoPro Bioscience, https://novoprolabs.com/tools/nls-signal-prediction) which applies a hidden Markov model in its algorithm (Nguyen Ba et al., 2009). See **Supplemental Table S1**.

### Phenotypic assays and microscopy of plants

Seedlings, trichomes (and EYFP-tagged proteins in trichomes), primary roots, and seeds were imaged with a Leica M125 fluorescence stereo microscope equipped with a GFP2 filter and a Leica DFC295 digital camera. Trichome quantification was performed as previously described (Khosla et al., 2014). For root hair quantification, vapor-sterilized seeds were sown on 1X MS medium (Murashige and Skoog, 1962) containing 0.8% Phytagel [w/v] (Sigma-Aldrich, P8169) and stratified at 4°C for 2-3 days. Seedlings were grown vertically under continuous light at 23°C for 2-3 days. To assess seed coat mucilage, seeds were stained with ruthenium red (Sigma-Aldrich, R2751) following a modified version of a published protocol (McFarlane et al., 2014). Approximately 20-25 seeds from each line were placed in 800 µl of 50 mM EDTA in a 24-well plate and incubated at room T with gentle shaking for 2 hr. Subsequently, the seeds were rinsed to remove the EDTA, and 800 µl of 0.01% ruthenium red [w/v] was added, followed by gentle shaking for 2 hr. The seeds were rinsed with dH2O (pH 7.5) and mounted on glass slides for imaging.

Live imaging of EYFP-tagged proteins in primary roots was performed with a Zeiss LSM 700 confocal microscope. Vapor-sterilized seeds were sown on 0.8% micropropagation agar [w/v] and stratified for 2-3 days at 4°C, followed by vertical growth at 23°C under continuous light. To visualize cell boundaries, seedlings were stained with 10 μg/ml propidium iodide (PI) for 2 min and rinsed with dH2O to remove excess stain. Seedlings were mounted in dH2O on a glass slide with a 25 x 25 mm coverslip. EYFP was excited at 488 nm and detected at 505-600 nm using a 40X oil immersion objective, while PI was excited at 555 nm and detected at 610-800 nm. The pinhole, gain, and laser intensities were adjusted accordingly to eliminate background fluorescence and to prevent photobleaching. Image processing was performed with ZEN (black edition) and Adobe Photoshop.

### Yeast 2-hybrid (Y2H) assay for GL2 dimerization

Full-length wild-type and mutant cDNAs in pENTR/D-TOPO vectors were transferred to pDEST32 bait and pDEST22 prey vectors (Proquest Two-Hybrid System, Invitrogen) using Gateway LR Clonase II Enzyme mix (ThermoFisher Scientific). Haploid yeast strains Y2HGold and Y187 were transformed with the pDEST32 and pDEST22 plasmids using LiAc/PEG transformation, and plated on -Leu and -Trp media (lacking leucine and tryptophan, respectively) to select for bait and prey plasmids. Yeast mating of Y2HGold and Y187, selection for diploids, and assaying for bait autoactivation and dimerization were performed as previously described (Mukherjee et al., 2022). Y2H results were imaged using a Gel Doc XR+ (Bio-Rad), after preparing four-fold serial dilutions from normalized cell cultures and spotting onto permissive and selective media using a 48-pin multiplex plating tool (‘Frogger’, Dankar, Inc.) followed by incubation at 30°C for 3 days. Protein expression of the baits was analyzed from liquid cell cultures expressing the pDEST32 plasmids. Total protein was extracted using the sodium hydroxide/trichloroacetic acid (NaOH/TCA) method and Western blotting was performed with a 1:500 dilution of GAL4 (DBD) RK5C1:sc 510-HRP mouse monoclonal antibody (sc-510HRP, Santa Cruz Biotechnology).

### EMSA

Halo-tagged wild-type PDF2 and mutant pdf2 proteins were produced from pIX-HALO plasmid DNAs using the TNT SP6 High-Yield Wheat Germ Protein Expression System as described previously (Mukherjee et al., 2022). Fluorescein amidite (FAM) labeled oligonucleotides, listed in **Supplemental Table S3**, were used to assess the DNA binding ability of the Halo-tagged proteins. The oligonucleotides were annealed according to a previously method (Mukherjee et al., 2022). In brief, 10 µl reactions consisted of 4 µl (100 μM) of oligonucleotides for a final concentration of 25 μM. To this, 6 µl annealing buffer was added, comprised of 100 mM Tris-Cl (pH 7.5), 1 M NaCl, and 10 mM EDTA. Annealing comprised of the following steps: 95°C for 2 min, 55°C for 5 min, a 90-min transition from 55° to 37°C, followed by a final hold at 4°C. For the EMSA, 20 µl reactions comprised of 12 µl of in vitro translated Halo-tagged proteins, 4 μl binding buffer (100 mM Tris-HCl (pH 7.5), 50 mM KCl, 5 mM DTT, 5 mM EDTA, 1.25 mg/mL salmon sperm DNA, 1.25 mg/mL BSA), 0.25 µl of 25 mM FAM labeled oligonucleotide, and 3.75 µl of dH2O (pH 7.5). The proteins were mixed with the binding buffer and incubated at 28°C for 10 min. Subsequently, 0.25 μl of the oligonucleotide and 3.75 µl of dH2O was added, and the mixture was incubated at 28°C for another 20- 25 min. 2 µl of 50% glycerol [w/v] was added to each sample prior to loading a 0.7% agarose [w/v] gel. Electrophoresis was conducted at 95 V for 1.5 hr at 4°C, followed by imaging of gels using a Typhoon Trio Imager (GE Healthcare) with the 520 nm emission filter for FAM (595 PMT voltage), 488 nm blue laser, and high sensitivity.

### GL2 NLS and IMPα structural prediction and protein-protein docking

To study the in silico protein-protein interaction between GL2 (NLS) and IMPα isoforms, we first predicted the 3D structure of the NLS peptide. This was achieved using PEP-FOLD (Lamiable et al., 2016), which utilizes computational methods to predict peptide structures for amino acid sequences. To predict 3D structures for IMPα isoforms, we performed homology modeling using MODELLER 9.21 (Webb and Sali, 2021) with the crystal structures of *Arabidopsis* IMPα-3 (PDB ID: 4TNM) and rice rIMPα-1 (PDB ID: 4B8J) as templates. The IMPβ Binding (IBB) domain, an autoinhibitory domain spanning the first ∼90 N-terminal residues, was removed prior to structural prediction. The IBB domain contains a segment of basic residues resembling an NLS, resulting in autoinhibition of the NLS interacting domain of IMPα in the absence of IMPβ (Cingolani et al., 1999; Chang et al., 2012; Jibiki et al., 2022). Following 3D structural prediction, loop refinement was performed to enhance the structural integrity and robustness of the interaction models. The model was evaluated using the SWISS-MODEL structure assessment tool (Waterhouse et al., 2018). To ensure stability and reduce entropy, energy minimization was achieved using MOLECULAR OPERATING ENVIRONMENT (MOE), an integrated computer-aided molecular design platform (Chemical Computing Group, Montreal, CA). Finally, the protein-peptide complex formed between the IMPα-3 and GL2 (NLS) was rendered using the High Ambiguity Driven protein-protein DOCKing (HADDOCK) web server (De Vries et al., 2010). The docked complexes were then analyzed using the Pymol and Discovery Studio (BIOVIA, version 2020) visualization tools to identify the interacting residues and bond types.

### Cytosolic yeast-2-hybrid (cyto-Y2H) assay and quantification of β-galactosidase activity

The in vivo protein-protein interaction between IMPα isoforms and wild-type GL2 versus gl2^ΔNLS^ was investigated using the split-ubiquitin-based Y2H system also known as a cyto-Y2H (Johnsson and Varshavsky, 1994). GL2 and gl2^ΔNLS^ were utilized as baits and were expressed as fusion proteins with an N-terminal OST4 anchor (Möckli et al., 2007). IMPα isoforms served as prey. Bait (pBT3-C-Ost4) and prey (pPR3-N) plasmids were transformed into NMY51 (*MATa his3*Δ*200 trp1-901 leu2-3, 112 ade2 LYS2*::*(lexAop)4-HIS3 ura3*::*(lexAop)8-lacZ ade2*::*(lexAop)8-ADE2 GAL4*) (Dualsystems Biotech) using standard LiAc transformation. Transformants were selected on minimal media lacking leucine and tryptophan (-Leu -Trp).

To assay interaction based on activation of the *lacZ* reporter gene, yeast cells containing bait and prey constructs were grown in 2 ml selective media (-Leu -Trp) in 24-well plates for 2 days. A volume of 0.5 ml was pelleted and subjected to freeze-thaw cycles to lyse the cells. After centrifugation, 30 µl of supernatant from each sample was transferred to a 96-well plate. For the enzyme assay, 80 µl of *ortho*-Nitrophenyl-β- galactoside (ONPG) was added to each sample, followed by incubation at 30°C. Reactions were terminated by addition of 1M Na2CO3 upon yellow color formation. A VictorV microplate reader (PerkinElmer) was used to measure OD405 of the reactions, and OD600 of the cell cultures prior to lysis. The β-galactosidase units were calculated according to the following formula: β-galactosidase Units = 1000 x OD405 / (Incubation Time x Culture Volume x OD600)

### Co-IP of recombinant proteins

The pGEX-6P-1 plasmids harboring GST-tagged IMPα isoforms were transformed into *E. coli* BL21 Rosetta 2 (DE3) (Novagen) cells. Single colonies were grown in 10 ml LB containing 100 µg/ml chloramphenicol and 100 µg/ml ampicillin on a shaker at 37°C. The next day, 3 ml cultures were inoculated into 300 ml 2xYT, followed by incubation at 37°C until the cells reached an OD600 of 0.6-0.7. Cultures were cooled on ice for 15 min, followed by induction with 0.1 mM IPTG for 18 hr at 18°C. After centrifugation at 4°C, cell pellets were frozen at -80°C. To prepare proteins, cells were sonicated in 10 ml of ice-cold lysis buffer (150 mM NaCl, 50 mM Tris, 1 mM DTT, 5% glycerol [w/v], 0.03% NP-40 [w/v], 5 mM MgCl2, 2% lysozyme [w/v]) supplemented with 1 mini-tablet of EDTA-free protease inhibitor (ThermoScientific, PIA32955) followed by high-speed centrifugation for 30 min at 4°C. Supernatants were centrifuged for another 25 min at 4°C. Protein concentrations in the lysates were estimated by Western blot. The TNT SP6 High-Yield Wheat Germ Protein Expression System (Promega, L3260) was used to generate Halo-tagged proteins starting with pIX-HALO plasmids as described previously (Mukherjee et al., 2022). A volume of 300 μl of GL2 bait (or mutant version) and 12 μl of IMPα prey proteins were mixed with 300 μl of binding buffer (150 mM NaCl, 50 mM Tris, 1 mM DTT, 5% glycerol [w/v], 0.02% NP-40 [w/v], and 5 mM MgCl2, and 1% BSA [w/v]). Binding reactions were incubated for 1 hr at room T with continuous rotation, followed by the addition of 15 μl of pre-washed glutathione agarose beads (ThermoScientific, PI16100). The samples were incubated for an additional 1 hr at room T with continuous rotation, and the beads were pelleted by centrifugation at 1500 x *g* for 1 min. The beads were washed five times, for 5 min each at room T, using a wash buffer (150 mM NaCl, 50 mM Tris, 1 mM DTT, 5% glycerol [w/v], 0.03% NP-40 [w/v], 5 mM MgCl2), with continuous end-to-end rotation. The bound proteins were eluted into 30 μl Laemmli buffer by heating at 99°C for 10 min. Samples were subjected to SDS-PAGE, followed by Western blotting using Anti-HaloTag monoclonal Ab (1:2000) (Promega, G9211) and THE GST monoclonal Ab (1:20000) (GenScript, A00865) primary antibodies, and Goat-Anti-Mouse IgG [HRP] (1:3000; GenScript, A00160) secondary antibody. Proteins were detected with SuperSignal West Femto Maximum Sensitivity Substrate (ThermoFisher Scientific) with an Azure c300 chemiluminescence imager (Azure Biosystems).

### Protein immunoprecipitation and protein digestion

Pull-downs were performed using EYFP-tagged GL2 as described (Wu et al., 2019). Protein from pulldowns were predigested for three hours with endoproteinase Lys-C (0.5 µg µL^-1^; Wako Chemicals, Neuss) at room temperature. After 4-fold dilution with 10 mM Tris-HCl (pH 8), samples were digested with Sequencing Grade Modified trypsin (0.5 µg µL^-1^; Promega) overnight at 37 °C. After overnight digestion, trifluoroacetic acid (TFA) was added (until pH ≤ 3) to stop digestion. Digested peptides were desalted over C18 tip (Rappsilber et al., 2003) and dissolved in 100 µl 80% acetonitrile and 0.1%.

### LC-MS/MS analysis of peptides and phosphopeptides

Tryptic peptide mixtures were analyzed by LC/MS/MS using nanoflow Easy-nLC1000 (Thermo Scientific) as an HPLC-system and a Quadrupole-Orbitrap hybrid mass spectrometer (Q-Exactive Plus, Thermo Scientific) as a mass analyzer. Peptides were eluted from a 75 μm x 50 cm C18 analytical column (PepMan, Thermo Scientific) on a linear gradient running from 4 to 64% acetonitrile in 240 min and sprayed directly into the Q-Exactive mass spectrometer. Proteins were identified by MS/MS using information-dependent acquisition of fragmentation spectra of multiple charged peptides. Up to twelve data-dependent MS/MS spectra were acquired for each full-scan spectrum acquired at 70,000 full-width half-maximum resolution. Fragment spectra were acquired at a resolution of 35,000. Overall cycle time was approximately one second.

Protein identification and ion intensity quantitation was carried out by MaxQuant version 2.2.0.0 (Cox and Mann, 2008) and the Andromeda matching algorithm (Cox et al., 2011). Spectra were matched against the *Arabidopsis* proteome (TAIR10, 35386 entries). Thereby, carbamidomethylation of cysteine was set as a fixed modification; oxidation of methionine was set as variable modifications. Mass tolerance for the database search was set to 20 ppm on full scans and 0.5 Da for fragment ions.

Multiplicity was set to 1. For label-free quantitation, retention time matching between runs was chosen within a time window of two minutes. Peptide false discovery rate (FDR) and protein FDR were set to 0.01, while site FDR was set to 0.05. Hits to contaminants (e.g., keratins) and reverse hits identified by MaxQuant were excluded from further analysis. The MaxQuant output protein groups.txt was used for further analysis. iBAQ-values were used (Schwannhäusser et al., 2009) for quantitative comparison.

## ACCESSION NUMBERS

The raw data from the proteomics experiment is deposited in ProteomeXChange under accession no. PXD044095. ANL2, AT4G00730; ATML1, AT4G21750; FWA, AT4G25530; GL2, AT1G79840; HDG1, AT3G61150; HDG2, AT1G05230; HDG3, AT2G32370; HDG4, AT4G17710; HDG5, AT5G46880; HDG7, AT5G52170; HDG8, AT3G03260; HDG9, AT5G17320; HDG10, AT1G34650; HDG11, AT1G73360; HDG12, AT1G17920; IMPA1, AT3G06720; IMPA2, AT4G16143; IMPA3, AT4G02150; IMPA4, AT1G09270; IMPA6, AT1G02690; PDF2, AT4G04890

## SUPPLEMENTAL DATA

Supplemental Figure S1. NLS positions and domain configurations of HD-Zip IV TFs from *Arabidopsis*.

Supplemental Figure S2. Conservation of NLS in HD-Zip IV TFs across the plant kingdom.

Supplemental Figure S3. The *gl2* NLS mutants are defective in seed mucilage production.

Supplemental Figure S4. Homodimerization is not impaired in *gl2* NLS mutants.

Supplemental Figure S5. NLS binding residues are conserved in IMPα isoforms from *Arabidopsis* and other eukaryotic organisms.

Supplemental Figure S6. Structural evaluation of IMPα-3 after model refinement and gap filling using homology modeling.

Supplemental Figure S7. Pull-down with EYFP:GL2 identifies proteins with functions in nuclear targeting.

Supplemental Figure S8. Tissue-specific expression of IMPα isoforms in *Arabidopsis*.

**Supplemental Table S1.** Bioinformatic prediction of monopartite NLS in HD-Zip IV and HD-Zip III TFs.

**Supplemental Table S2.** IMPα-3 and GL2 NLS interaction report from computational docking.

**Supplemental Table S3.** Oligonucleotides used in this study.

**Supplemental Data Set S1.** Protein list for the interactome of GL2. (in xls file)

## ACKNOWLEDGEMENTS

We thank Mareike Heilmann and Jae Bok Heo for their gifts of plasmids and seeds, respectively, Lin Xi for help with the pull-down experiment, and Endymion Cooper and Charles Delwiche for providing unpublished assemblies of *Spirogyra pratensis* homologs. Kyle A. Thompson aided in generating the NLS deletion in *GL2*. We are grateful for Joel Sanneman and the Confocal Core Facility at the College of Veterinary Medicine for assistance.

## AUTHOR CONTRIBUTIONS

B.A., R.L.-R. and K.S. conceived of and designed the experiments. B.A. and K.S. wrote the manuscript. R.L.-R., T.M. and B.A. created plant constructs and transgenic lines. B.A. and J.R.C. performed molecular docking simulations. K.S. and W.S. performed proteomics and mass spectrometry. H.V.N. conducted trichome and root hair quantification as well as ruthenium red staining of seeds. A.L.W. and B.A. performed the cytosolic Y2H experiments and quantitative assays. T.M. performed classical Y2H experiments to assay homodimerization. B.A. performed confocal microscopy, EMSA, Co-IP, and all other phenotypic and molecular analyses.

